# Coronavirus membrane protein with a fluorescent protein tag enables tracking of virus particles in live cells

**DOI:** 10.64898/2025.12.29.696688

**Authors:** Bereket Stefanos, Noe R. B. Green, Isha Nadig, Lynn A. Lapierre, James R. Goldenring, Ian B. Hogue, Honor L. Glenn, Brenda G. Hogue

## Abstract

Coronavirus particles assemble at endoplasmic reticulum Golgi intermediate compartment (ERGIC) membranes and exit from host cells via secretory organelles that are not well defined. The interplay between viral components and intracellular transport pathways that facilitate assembly and egress are not fully understood and recent studies suggest that multiple pathways maybe involved. Reverse genetics was used to develop a model system to further understand the assembly and egress processes. Mouse hepatitis coronavirus (MHV-A59) was genetically engineered to express the membrane (M) protein fused to green fluorescent protein (M-GFP), with the chimeric gene cloned in place of the open reading frame (ORF) 4 coding region in the RNA genome. The recovered M-GFP virus also expresses wild-type (WT) M protein (M WT) from its native ORF. The M-GFP virus exhibited morphology and growth properties like WT virus. M-GFP and WT M proteins colocalized early in infection, but less M-GFP trafficked toward the cell surface at later times, suggesting that the fusion protein is incorporated less efficiently into virus particles. M-GFP was stably expressed through at least four virus passages. Early passage virus M-GFP virus particles were visualized by confocal and Total Internal Reflection Fluorescence (TIRF) microscopy in live cells. The fluorescently labeled virus particles represent a new tool for coronavirus intracellular trafficking and egress studies. Such studies can also help provide a more detailed understanding of infection and disease processes to provide new insight for development of new therapeutic strategies.

**IMPORTANCE:** Coronavirus assembly, intracellular transport, egress, and the virus-host interactions involved in these processes are not fully understood. The M protein is the most abundant virion structural component, which forms the scaffold for assembly of the viral envelope. It facilitates incorporation of the viral nucleocapsid and the other viral membrane proteins, spike (S) and envelope (E). In this study we engineered M protein tagged with GFP, which is expressed along with wild-type (WT) M from the viral genome. M-GFP interacted with WT M protein and co-assembled into virus particles. The resulting fluorescently labeled coronavirus particles open opportunities to use state of the art microscopy to monitor virus assembly, trafficking, and egress in live cells to facilitate deeper understanding of the molecular and cell biology of these processes. The ability to visualize particles also opens opportunities to help further understanding of steps in viral replication and pathogenesis for potential therapeutic targets.

## INTRODUCTION

The *Coronaviridae* family consists of enveloped, positive-sense, single-stranded RNA viruses that infect humans and a wide range of animals, including domesticated and wild mammals and avian species (1, 2). The significance of highly pathogenic human coronaviruses (hCoVs) emergence was clearly shown during the recent SARS-CoV-2 COVID-19 pandemic. Even though the pandemic phase is over, SARS-CoV-2 variants continue to emerge as the virus appears to now be endemic in the human population. Highly pathogenic viruses are likely to arise again, given that three new hCoVs have emerged in the past two decades. Thus, it is important to continue increasing our understanding of the molecular and cell biology of these viruses, to provide fundamental knowledge that will be needed for development of new therapeutics and vaccines targeting future emerging viruses.

Coronaviruses assemble and bud at intracellular membranes of the endoplasmic reticulum Golgi intermediate compartment (ERGIC) (3, 4). All coronaviruses express three membrane structural proteins: spike (S) that binds host receptors, membrane (M) the major structural scaffold for the envelope, and small envelope (E) that is a viroporin. The non-membrane bound nucleocapsid (N) phosphoprotein encapsidates the genomic RNA to form what has long been described as a helical nucleocapsid packaged inside virus particles. Recent studies suggest that, rather than a helical structure, the N protein interacts with the genomic RNA as beads-on-a-string (5, 6). The nucleocapsid with the RNA interacts with the membrane proteins to drive its envelopment. The M protein, the focus of this report, is the most abundant protein in the viral envelope and plays key roles in virus assembly, as it forms the scaffold on which the envelope is formed. Multiple studies have also provided support for a role of the M protein in RNA packaging (7-10).

Coronavirus M proteins have a short amino terminus domain located outside the virion, followed by three transmembrane (TM) domains, and a long (∼100 amino acid) carboxy tail extending into the interior of particles (reviewed in (11)). The M protein organizes the assembly process by participating in M-M as well as M-S and M-N protein interactions (12-17). For most coronaviruses, co-expression of the M and E proteins alone drives virus-like particle (VLP) assembly (18, 19). The structure of SARS-CoV-2 M protein was recently determined by cryo-electron microscopy (cryo-EM) (20, 21). The protein forms homodimers that exist in two conformational states, described as the long and short forms (22).

To facilitate studies directed at increasing our understanding of viral protein and viral-host protein interactions during assembly and egress, we tagged M protein with GFP (M-GFP) at the carboxy end of the protein. The fusion protein was engineered into the MHV genome for co-expression with WT M. M-GFP is incorporated into virus particles, and our results demonstrate that it can be used to track virus particles in live cells. This tagged virus opens new opportunities for in-depth studies of virus assembly and trafficking, replication and pathogenesis.

## RESULTS

### Construction of a recombinant MHV expressing M-GFP fusion protein

Plasmid-based reverse genetics was used to engineer fluorescently tagged MHV particles to allow real-time tracking in cells (Fig. 1B). The coding sequence for green fluorescent protein (GFP) was appended to the carboxy end of the MHV M gene, with an intervening linker sequence consisting of small polar residues. AlphaFold2 modeling of the M-GFP protein predicts that the overall structures for M or GFP are not significantly altered in the fusion protein (Fig. 1C). The M-GFP fusion sequence was cloned into open-reading frame (ORF 4) in the MHV genome (Fig. 1A). The WT M gene was retained in normal position in the genome for expression along with M-GFP.

**Fig 1.**
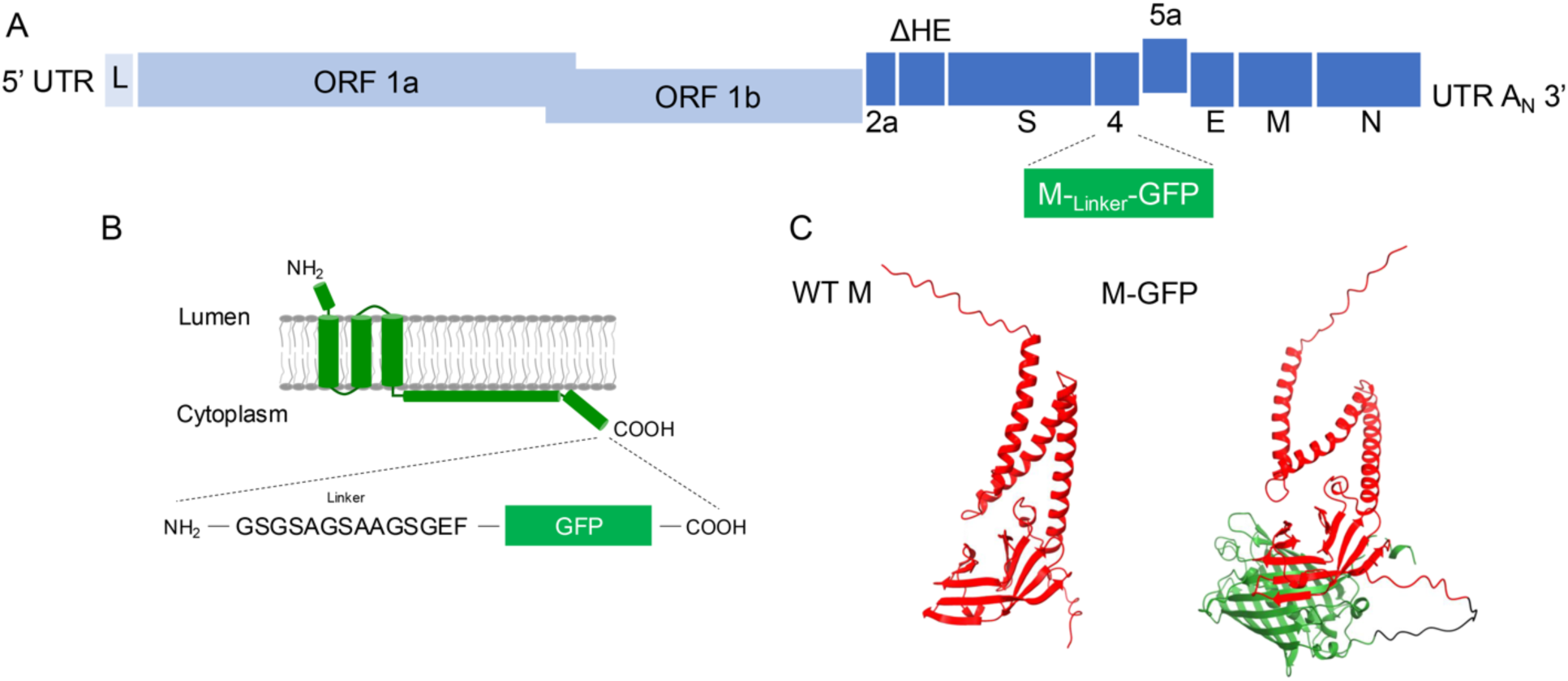
M protein fused to GFP was inserted into the MHV genome. (**A**) The M-GFP fusion sequence was inserted into ORF4. (**B**) When the M-GFP fusion protein is synthesized in the ER, the linker sequence and GFP are predicted to be on the cytoplasmic side of the membrane. (**C**) AlphFold2 prediction indicates that addition of GFP does not alter the folding of M.

### Recovery of a M-GFP fluorescently labeled virus

Virus containing untagged WT M and M-GFP was recovered, and the presence of M-GFP fusion gene in ORF 4 of genomic RNA was confirmed by sequencing. The isolated virus was plaque purified three times, and the presence of the fusion gene was reconfirmed by sequencing prior to generating working stocks for further experiments. Parental WT MHV and MHV M-GFP viruses were grown in mouse 17Cl1 cells. Extracellular viral supernatants were clarified before purification on sucrose gradients, followed by pelleting through a sucrose cushion. Undiluted virus suspensions from the purified samples were dropped directly onto a glass coverslip and imaged immediately by confocal microscopy. The M-GFP virus samples produced strong, green, tightly packed puncta at or below the resolution limit of light microscopy (Fig. 2A). The WT MHV sample had no discernible fluorescence when imaged under identical conditions. This indicated that GFP-tagged M protein is incorporated into virus particles that match the density of WT MHV, and that the fusion protein fluoresces as expected for functional GFP.

**Fig 2.**
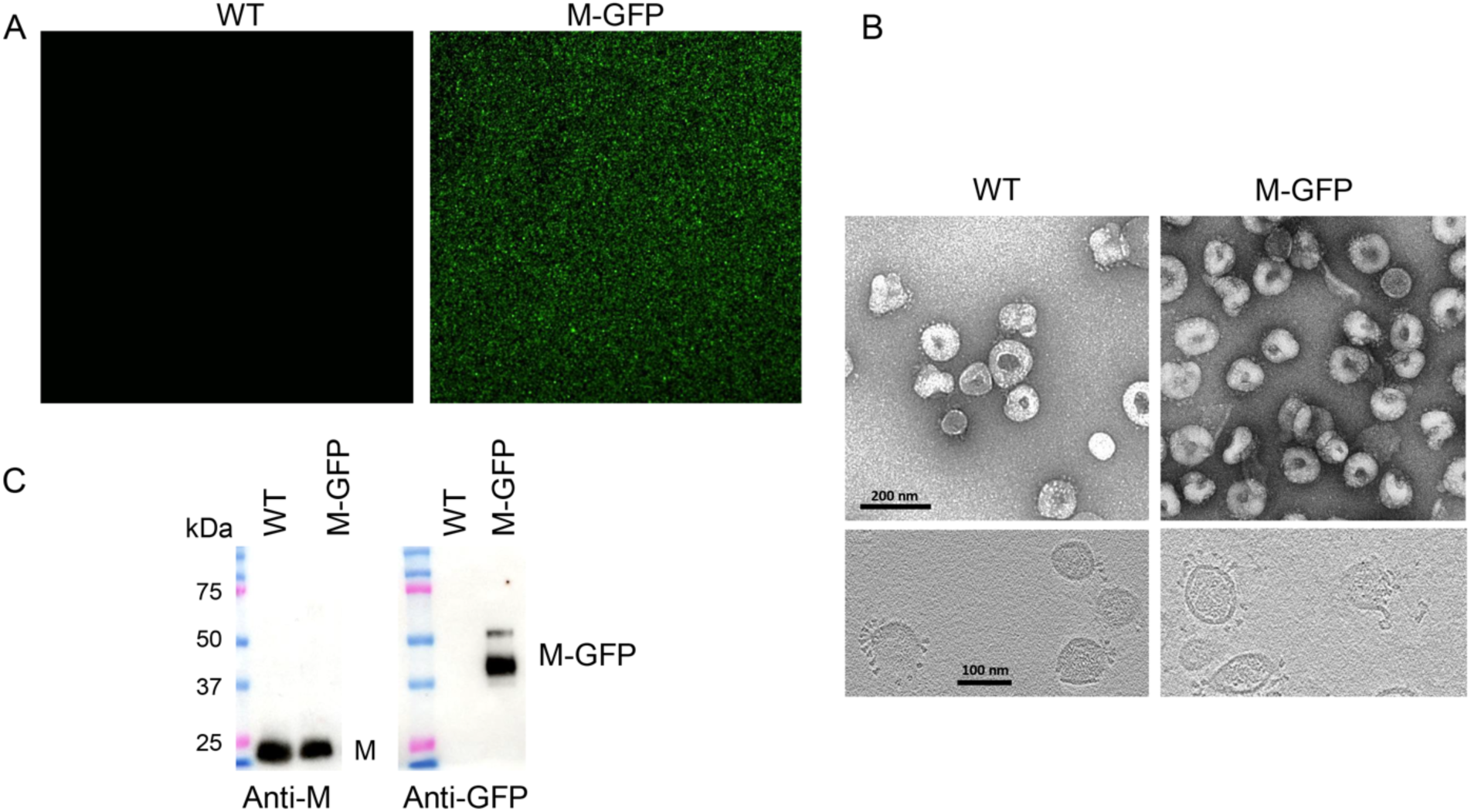
M-GFP MHV particles are fluorescent and structurally like WT MHV. (**A**) WT MHV and M-GFP MHV were purified on a sucrose gradient and fractions containing virus were pelleted through a sucrose cushion. Suspensions were diluted approximately 1:5, dropped onto a coverslip and imaged in the GFP channel by confocal microscopy. (B) Aliquots of purified virus were negatively stained and imaged by EM (top panels) or plunge frozen and imaged by cryo-EM (bottom panels). (C) Purified virions were analyzed by SDS-PAGE and western blots probed with antibodies against M or GFP.

Virus particles from the same purified preparations described above were also imaged by electron microscopy. Aliquots of the pelleted virus suspensions were spotted onto formvar/carbon coated copper grids and virions were imaged by transmission electron microscopy (TEM) with negative staining (Fig. 2B, top images). Samples were also vitrified by plunge freezing and imaged by cryo-EM (Fig. 2B, bottom images). Like WT MHV particles, M-GFP particles were roughly spherical with spike trimer projections, and were 80-120 nm diameter, typical for coronavirus particles (23-25).

### Characterization of MHV M-GFP virus

Western blotting was used to further confirm M-GFP assembly into virus particles. Equivalent amounts (1 x 10^7^ pfu) of the gradient purified particles were lysed and analyzed by SDS-PAGE and western blotting. Blots were probed with antibodies against the M or GFP proteins (Fig. 2C). The WT M protein was strongly detected with the anti-M antibody for both WT and M-GFP particles (left panel). M-GFP was present only in the M-GFP virus particles (right panel), further confirming assembly of the fusion protein into virion particles.

The predicted molecular weight of MHV M is 26 kDa without O-linked glycosylation at two sites on the amino end of the protein. Coronavirus M proteins characteristically migrate slightly faster than the calculated molecular weight under denaturing conditions during electrophoresis, as seen in Fig. 2C. The predicted molecular weight for M-GFP with the linker is ∼54 kDa. Anti-GFP recognized two bands. One migrated between the 37 and 50 kDa markers and a second one slightly above 50 kDa. The hydrophobic nature of the M protein and/or the net negative charge under SDS-PAGE running conditions likely accounts for the faster mobilities observed for both M and M-GFP proteins.

To further analyze the two M-GFP protein species, and to exclude the possibility that the smaller ∼37 kDa M-GFP band represents a cleaved or truncated protein, two plasmids were constructed for M-GFP expression with a V5 epitope tag at either the amino or carboxy end of the fusion protein. The plasmids were transfected into 293T cells and cytoplasmic lysates were analyzed by western blotting. The ∼37 kDa and ∼50 kDa protein bands were both recognized by the V5 specific antibody, when expressed from the amino or carboxy-tagged constructs, indicating that both protein bands represent full-length M-GFP (Fig. S2A). The V5 tagged M-GFP proteins were also expressed in mouse L2 cells. Protein expression was analyzed using antibodies against the amino terminus of WT M or the V5 tag and imaged by confocal microscopy at 24 h after transfection. The V5 tag at either the amino end or carboxy end of the M-GFP fusion protein were each detected. The signals for V5, WT M and GFP colocalized (Fig. S2B), further supporting the presence of intact ends on the fusion protein.

We expected that M-GFP would be incorporated into virions through M-M interactions with the WT M protein. In this case, we reasoned that the proteins could be co-immunoprecipitated. To investigate this, mouse 17Cl1 cells were infected with WT or M-GFP viruses. Intracellular and extracellular fractions were harvested at 24 hpi. Virions in the extracellular fraction were pelleted through a sucrose cushion. Intracellular and extracellular lysates were divided into three aliquots. One part was used as the non-immunoprecipitated cell lysate (input), the second was immunoprecipitated with anti-M J1.3 antibody that recognizes the amino end of the protein (26). The third part was immunoprecipitated with a GFP-specific monoclonal antibody (Invitrogen, GF28R). Proteins eluted following immunoprecipitation were run on parallel SDS-PAGE gels. The input samples were included for direct western blotting. Blots were probed with polyclonal anti-M 9246 or GFP polyclonal rabbit antibodies (Fig. 3). WT M and M-GFP were strongly detected in the input samples for both WT and M-GFP intracellular and extracellular fractions (Fig. 3A-B, lanes 1-3). The J1.3 M antibody immunoprecipitated WT M from both intracellular lysates and extracellular virions from cells infected with either WT (Fig. 3A, lane 5) or M-GFP virus (Fig. 3A, lane 6). M-GFP was co-immunoprecipitated with WT M from M-GFP infected cells and virions (Fig. 3B, lane 6). Reciprocally, the GFP antibody immunoprecipitated M-GFP (Fig. 3B, lane 9) and co-immunoprecipitated WT M (Fig. 3A, lane 9). This strongly indicates that M-GFP interacts with WT M, suggesting that even if M-GFP itself is partially defective in function, it is nevertheless able to be incorporated into virions via interactions with WT M.

**Fig 3.**
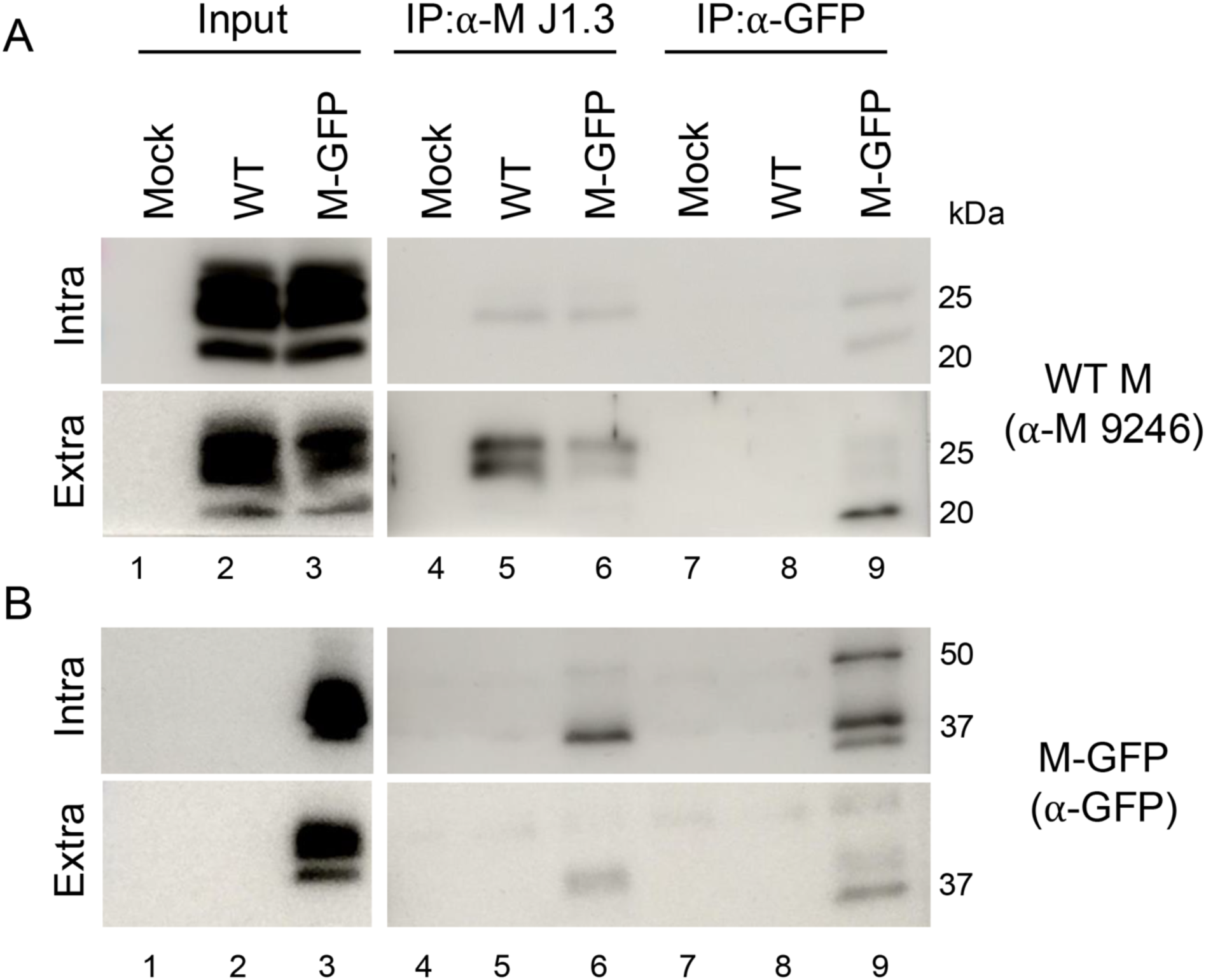
M-GFP and WT M proteins Co-IP. L2 cells were uninfected or infected with the WT virus or the M-GFP virus. Infected and uninfected intracellular and extracellular factions were lysed in a RIPA buffer and the M protein or M-GFP protein were immunoprecipitated (IP) out with an amino targeted M antibody and a GFP antibody respectively followed by a pull down with Sepharose A beads. IPed lysates were run on a western probing for the WT M protein (A) and the M-GFP protein (B). Lanes 1-3 are the input controls for both mock and infected cells. Lanes 4-6 are blots for the WT M immunoprecipitation with the α-M J1.3 antibody and lanes 7-9 are blots for the M-GFP immunoprecipitation with the α-GFP antibody. The α-M J1.3 pulls down the WT M protein in WT M infected and M-GFP infected cells in both the extracellular and intracellular fractions (A lanes 5 and 6). The α-M J1.3 also pulls down the M-GFP protein in the M-GFP infected cells in both the extracellular and intracellular fractions (B lanes 5 and 6). The α-GFP pulls down the WT M proteins and the M-GFP protein in the extracellular and intracellular lysate of M-GFP infected cells (A/B lane 9)

### M-GFP virus growth properties

To further characterize the M-GFP recombinant virus the growth properties were compared with WT virus. Mouse 17Cl1 cells were infected in parallel with WT or M-GFP at a multiplicity of infection (MOI) of 0.01. The multi-step growth kinetics showed that the M-GFP virus grew at a similar rate, with no statistically significant difference from WT MHV at the final time point of 48 hpi (p=0.06) (Fig. 4A). The recombinant virus displayed a plaque phenotype like the WT virus, but also included some smaller plaques, which were always present even after multiple rounds of plaque purification (Fig. 4B).

**Fig 4.**
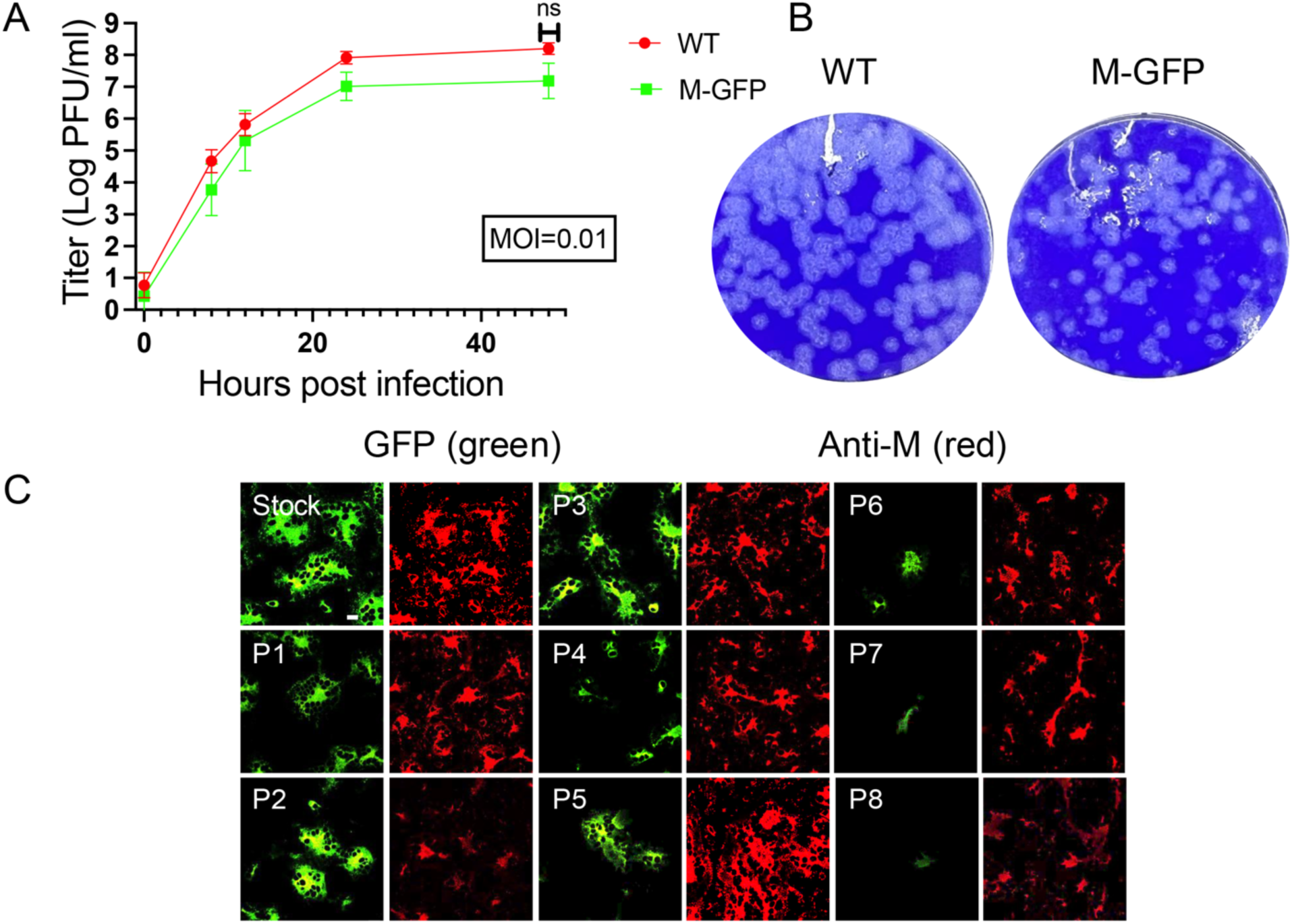
M-GFP exhibit growth properties like WT MHV. (A) M-GFP and WT MHV were plaqued in parallel in L2 cells. M-GFP virus displays plaque morphology like WT and smaller plaques present after multiple rounds of plaque purification. Growth kinetics of the M-GFP virus was monitored in comparison to the WT virus (B). (B) 17Cl1 cells were infected at a MOI of 0.01 and titers were measured by plaque assay at the indicated times. M-GFP virus growth kinetics were like WT MHV, with no significant titer difference at 48 hpi, p=0.0676. (C) The stability of the M-GFP protein in the M-GFP virus was monitored over 8 passages in L2 cells (C). Direct visualization of M-GFP showed loss of the protein at passage 5 while maintaining WT M expression visualized through IF. Scale bar = 20 µm

The stability of the M-GFP virus was monitored through eight passages (P8) in mouse L2 cells. Cells were imaged at each passage detecting GFP fluorescence. Cells were also fixed and M antibody 9246 was used to monitor expression of WT M. GFP expression was abundant in the earlier passages, including the stock virus, P1, P2, and P3 (Fig. 4C). However, by P4-5, the GFP signal declined and by P8 there was very little GFP expression. The level of WT M expression was maintained throughout all passages, indicating no major changes in infection efficiency. The P8 M-GFP virus was sequenced, and it showed that the entire M-GFP protein coding region was lost. These results indicate that the virus is under selective pressure to lose M-GFP expression from ORF4. Multiple attempts to replace the WT M with M-GFP were not successful, indicating that while M-GFP can be assembled into virus particles, WT M is also required for full viral assembly.

### M-GFP traffics similarly to WT M early in infection

Early in infection, 4-8 hpi, M and other structural proteins localize to the ERGIC/Golgi region, where viral assembly occurs (3, 27). Between 8 and 12 hpi, newly assembled virus particles begin trafficking to the cell periphery, whereas most unassembled structural proteins are retained at the ERGIC/Golgi region (3, 27). During the 8-12 hpi timeframe, some S protein traffics separately from virus particles to the cell surface where it mediates cell fusion resulting in multinucleate syncytia.

To determine whether the M-GFP virus recapitulates these established protein localization and trafficking patterns, L2 cells were infected with M-GFP virus at MOI=5, fixed and immunolabeled at 4, 6, and 8 hpi, and imaged by confocal microscopy. At the earliest time point, WT M and M-GFP strongly colocalize at ERGIC membranes (Fig. 5, 4 hpi, arrows). Beginning at 6 hpi, and more prominently at 8 hpi, discrete puncta containing WT M and M-GFP are apparent outside of the ERGIC region, consistent with virus particle assembly and trafficking (Fig. 5, 6-8 hpi). However, puncta at the cell periphery appear to contain more WT M, whereas more M-GFP is retained at the ERGIC. Similar results were obtained in mouse 17Cl1 and human A549 cells expressing the MHV cell surface receptor, CEACAM-1 (data not shown).

**Fig 5.**
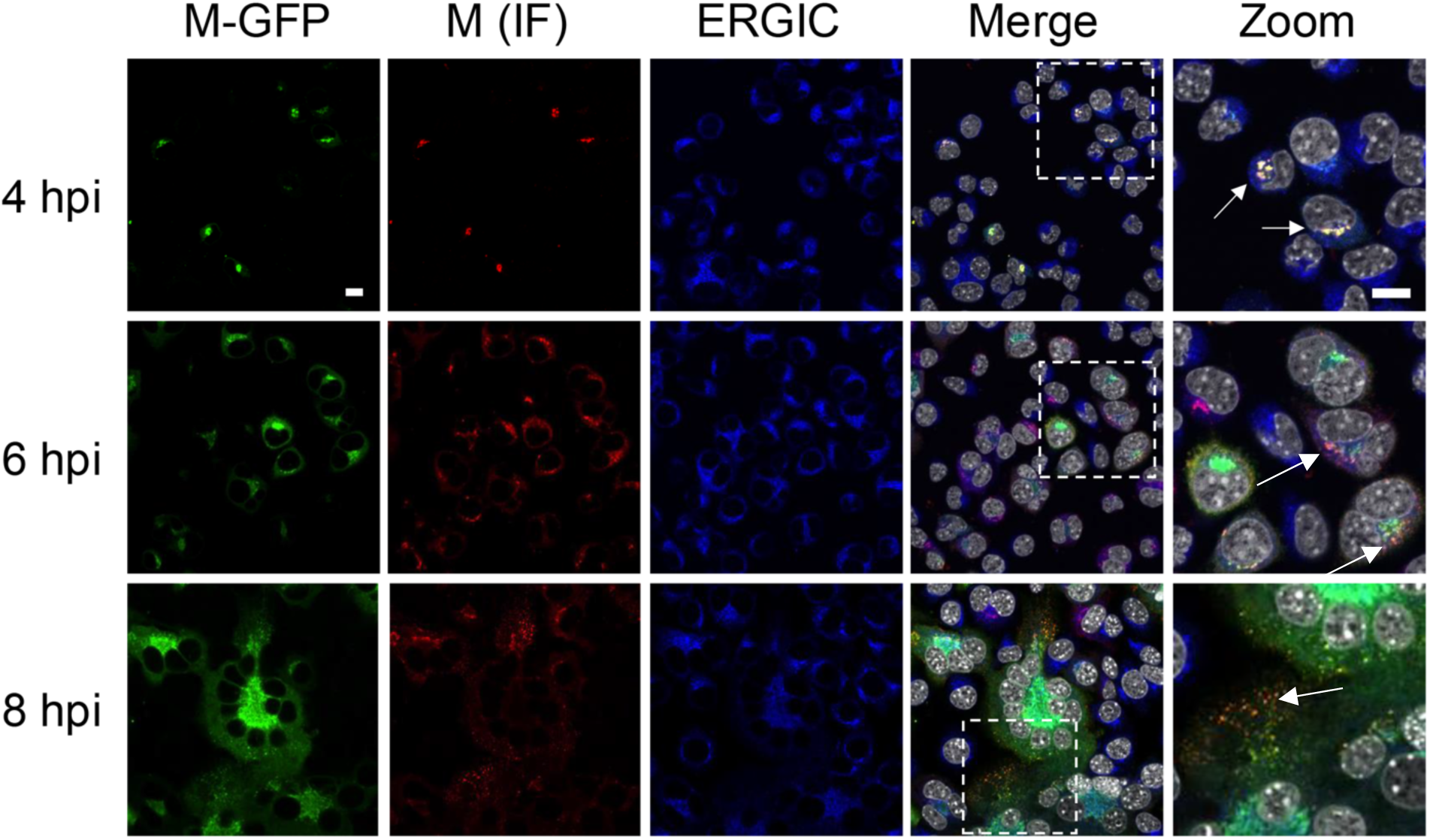
M-GFP traffics with WT M during MHV infection. L2 cells were fixed at 4, 6, and 8 hpi with M-GFP MHV, then immunolabeled for WT M and ERGIC. Nuclei were labeled with DAPI. Samples were imaged by confocal microscopy. Zoomed regions are indicated by the dashed square in the merged image. Green=M-GFP; red=WT M; blue=ERGIC, gray=nuclei. Images represent typical results from at least three fields imaged from each of three, independent experiments. Scale bar = 10 µm.

To quantify these observations, we performed colocalization analysis. Manders’ coefficients (M1 and M2) are intensity-independent measures of the fraction of one fluorescent signal that overlaps with another (and vice versa). Pairwise colocalization images from the representative 6 hpi timepoint (Fig. 5) are shown in Fig. 6A. At 4 hpi, a large majority of WT M and M-GFP (M1 ∼0.8) overlapped with the ERGIC marker (Fig. 6A). As infection progressed, the fractions of WT M and M-GFP overlapping the ERGIC decreased significantly, by about 50% by 8 hpi (p< 0.0001), indicating that both protein species traffic out of this compartment (Fig. 6B). WT M and M-GFP followed similar and parallel courses in the time dependent change in ERGIC colocalization. Pairwise overlap of M-GFP and WT M with the ERGIC compartment at 6 hpi are shown in the second and third panels respectively in Fig 6A. These results are consistent with both WT M and M-GFP being incorporated into virus particles and trafficking to the cell periphery. The fraction of M-GFP signal that overlapped with WT M (M1), was about 0.73 across time points (Fig. 6C). This indicates that most M-GFP colocalizes with WT M. The corresponding fraction of WT M that overlaps with M-GFP (M2) was significantly lower (p<0.0001), at about 0.53 (Fig. 5D). While we observed a consistent downward trend in the fraction of WT M overlapping with M-GFP over time (M2), this decrease over time was not statistically significant. The pairwise overlap of WT M and M-GFP at 6 hpi is shown in the left panel of Fig 6A. We also analyzed the colocalization of WT M and M-GFP by Pearson’s correlation, an intensity dependent measure. WT M and M-GFP were positively correlated, and while this correlation also had a downward trend, this decrease over time was not statistically significant (p=0.07) (Fig. 6E). This indicates that the intensity of WT M and M-GFP have a positive correlation across all pixels at all measured time points.

**Fig 6.**
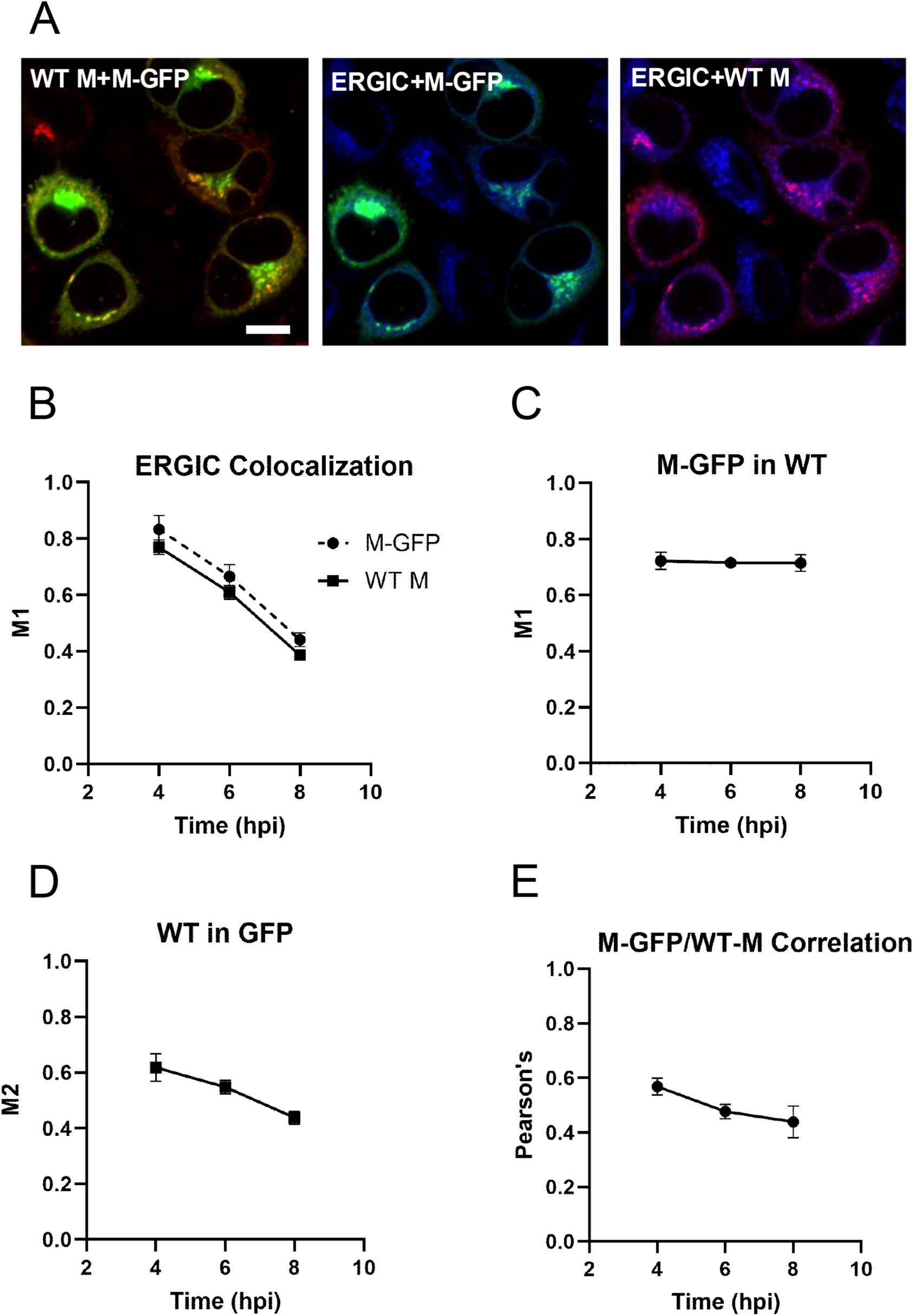
Quantitative analysis of M-GFP and WT M intracellular localization. (**A**) Pairwise representation of WT M (red) and M-GFP (green), left panel; ERGIC (blue) and M-GFP (green), middle panel; ERGIC (blue) and WT M (red), right panel. This is the same field shown in Fig 5, 6 hpi Zoom. Scale bar = 10 µm. (**B**) Mander’s correlation coefficients (M1) were calculated for both the fraction of WT M signal and fraction of M-GFP signal that colocalized with the ERGIC at 4, 6, and 8 hpi. Two-way ANOVA revealed that there was a time-dependent loss of both species from the ERGIC over the time frame analyzed (p<0.0001) and that slightly more M-GFP localized to this compartment relative to WT M (p=0.03). Error bars indicate SEM. Data were pooled from at least two representative fields imaged from each of three, independent experiments. (**C**) Mander’s correlation coefficient was calculated for the fraction of M-GFP signal that colocalized with WT M signal at 4, 6, and 8 hpi (M1). ANOVA revealed no significant time dependent change in colocalization. Error bars indicate SEM. Data were pooled from at least three representative fields imaged from each of three, independent experiments. (**D**) Mander’s correlation coefficient was calculated for the fraction of WT M signal that colocalized with M-GFP at 4, 6, and 8 hpi (M2). ANOVA revealed no significant time dependent change in colocalization. Error bars indicate SEM. Data were pooled from at least three representative fields imaged from each of three, independent experiments. (**E**) Pearson’s correlation coefficient indicates a positive correlation in intensity for M-GFP and WT M signal at 4, 6, and 8 hpi. ANOVA revealed no significant time dependent change in correlation. Error bars indicate SEM. Data were pooled from at least three representative fields imaged from each of three, independent experiments.

These results are consistent with the conclusion that M-GFP localizes and traffics similarly to WT M. However, our qualitative observations and the trends in colocalization measurements (albeit not statistically significant) suggest that M-GFP may be less efficiently incorporated into virus particles, compared to WT M. Thus, a greater fraction of M-GFP remains localized to the perinuclear ERGIC region, whereas WT M is more efficiently incorporated into virus particles and trafficked to the cell periphery. Nevertheless, M-GFP is incorporated into virus particles (Fig. 2-3) and trafficked to the cell periphery (Fig. 5A, 6A), making it a useful marker of virus particle trafficking and egress.

### Live cell tracking of M-GFP

Our data suggest that M-GFP may be incorporated into mature virus particles less efficiently than WT M. This led us to question whether the GFP tagged protein was sufficiently abundant at the cell surface to serve as a marker for viral trafficking near the spatial-temporal point of viral egress from the cell. To address this question, we used live-cell Total Internal Reflection Fluorescence (TIRF) microscopy. This approach allows visualization of fluorophores within a highly limited depth of field, less than 200 nm from the bottom surface of cells adhered to a standard cover glass, while excluding fluorescence from deeper in the cytoplasm. We infected 17Cl1 cells with M-GFP MHV and imaged the live cells at 6-7 hpi. Time-lapse TIRF imaging revealed many green-fluorescent puncta with apparent diameters of approximately 480 nm, consistent with diffraction-limited virus particles, close to the cell surface. Some of these structures were stationary, while others moved rapidly (Fig. 7A, and Supplement S3). We used the Nikon Elements software for post-processing, and the Elements tracking module to track and analyze the moving particles. We successfully tracked moving particles over an average path length of 6.67 µm before particles stopped or moved out of the imaging plane. Particles exhibited an average velocity of 1.67 µm/s (Fig. 7B, Table 1, and Supplement S4), consistent with the typical velocities of microtubule-based motors (28, 29). Thus, moving puncta likely represent virus particles in the lumen of a secretory organelle undergoing microtubule-based transport prior to exocytosis. These results show that the M-GFP virus has significant utility for tracking viral transport at and near to the plasma membrane of infected cells.

**Fig 7.**
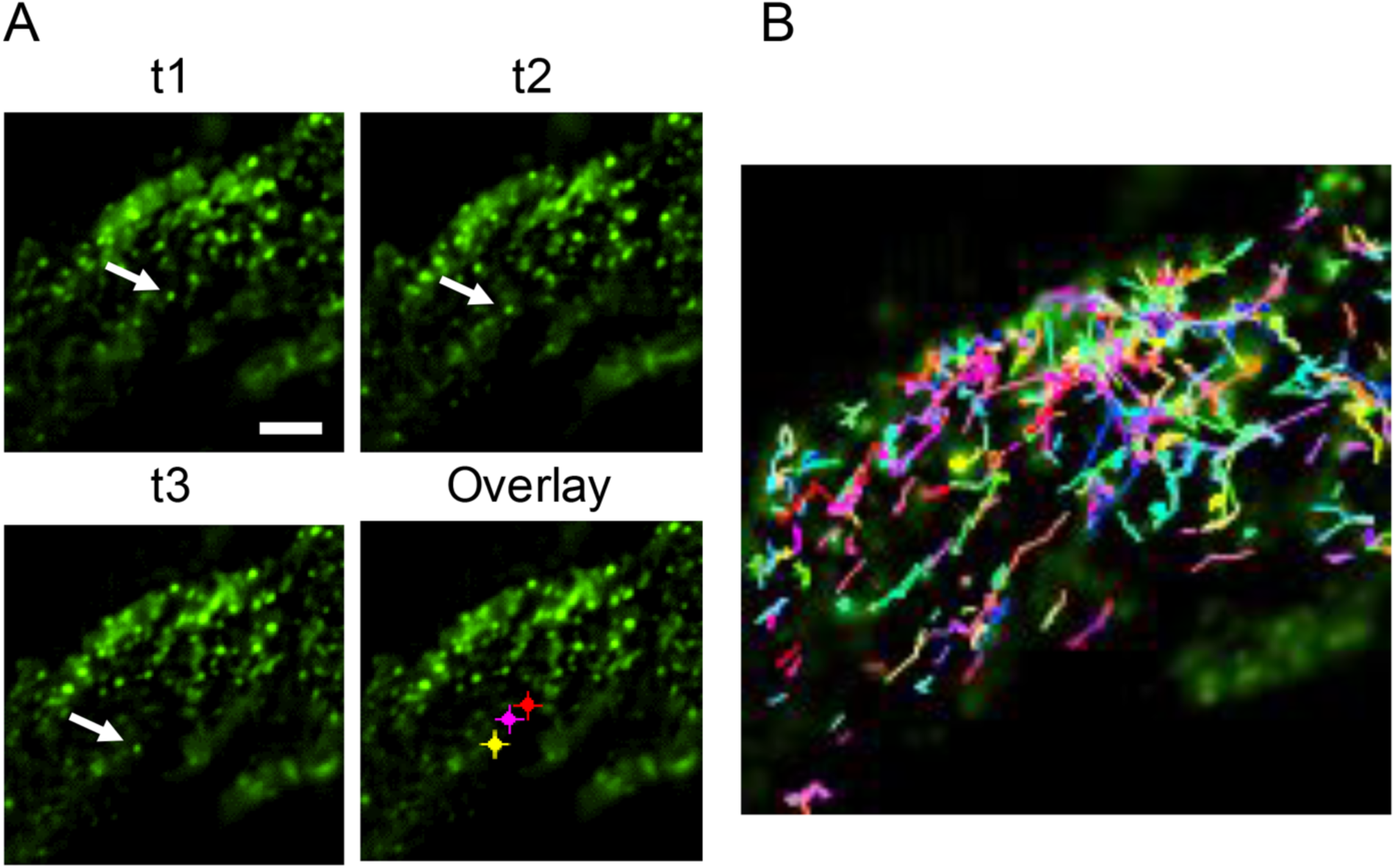
Tracking of intracellular M-GFP by Total Internal Reflection Fluorescence (TIRF) microscopy. (**A**) 17 Clone 1 (17C1) cells were infected with M-GFP MHV then imaged live by TIRF microscopy at 6-7hpi. Timelapse images were collected for up to 2.5 minutes per field. A cropped field of view is shown with three time points (t1, t2, t3) spanning ∼5.5 seconds with the position of one GFP-positive particle indicated (arrow). Overlay image shows the t3 time point image with each position Indicated. Red=t1; magenta=t2; yellow=t3. Scale bar = 5 µm. (**B**) Particle tracking was done on the same field as A. Tracks of moving particles are shown for ∼60 seconds of real time data collection.

**Table 1.**

|  | Length<br>[μm] | Duration<br>[sec] | Speed<br>[μm/sec] | Eq<br>Diameter<br>[μm] |
| --- | --- | --- | --- | --- |
| Mean | 6.57 | 6.16 | 1.67 | 0.48 |
| SEM | 0.32 | 0.99 | 0.25 | 0.01 |
| Length = moving particle total pathlength<br>Duration = particle motion time recorded<br>Speed = speed along path<br>Data were pooled from at least four fields from two independent experiments. |  |  |  |  |

### M-GFP puncta at the cell periphery colocalize with Spike (S) protein

While some viral structural proteins, like S and N, may traffic within the cell prior to or independent of virus particle assembly, M and E proteins are mainly retained at the ERGIC until assembled into virus particles (30). Accordingly, WT M and M-GFP are observed at the ERGIC at earlier times post-infection, before trafficking to the periphery as discrete puncta at later times (Fig. 5 and Fig. 6). To determine whether the M-GFP puncta at the cell periphery represent intact viral particles, we infected 17Cl1 cells with M-GFP virus, fixed and immunostained cells at 6.5 hpi, then imaged by TIRF microscopy. Consistently, M-GFP colocalized with WT M and S, detected by immunofluorescence, in discrete puncta (Fig. 8). Thus, M-GFP puncta near the cell surface likely represent mature virus particles.

**Fig 8.**
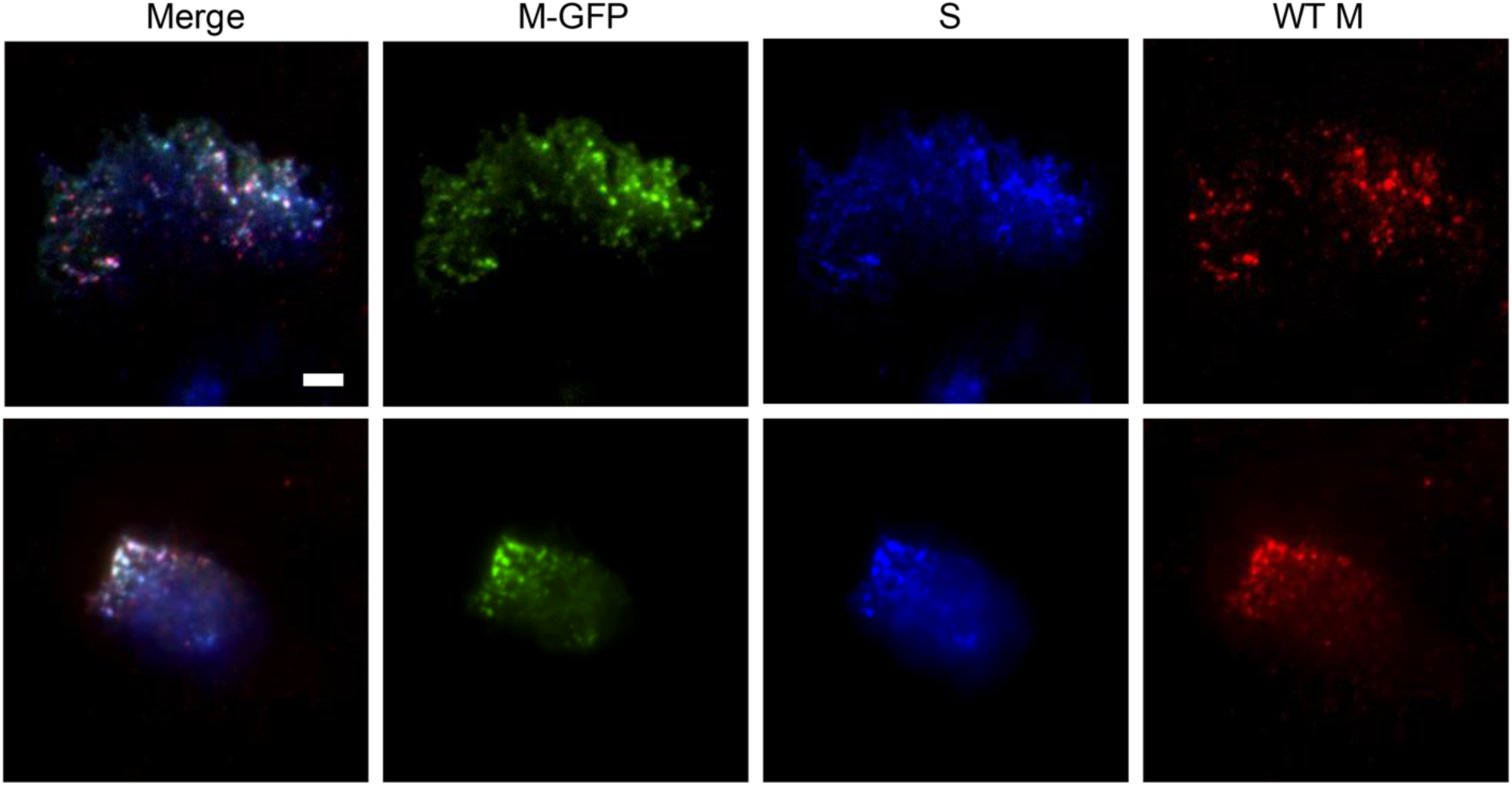
M-GFP colocalizes with WT M and Spike protein (S). 17C1 cells were infected with M-GFP MHV then fixed at 6.5 hpi and immunolabeled for WT M and Spike protein. Three color TIRF was used to show colocalization of M-GFP (green), WT M (red), and S (blue) at the plasma membrane. Two typical fields are shown from at least 4 images each from two independent experiments. Scale bar = 5 µm.

## DISCUSSION

We have demonstrated, to our knowledge for the first time, a coronavirus expressing a functional, fluorescent protein-tagged M protein that is assembled into virus particles that can be tracked in live cells. We chose to tag the M protein because it is strongly retained at the ERGIC prior to assembly and traffics to the cell periphery mainly following incorporation into virus particles. Thus, M-GFP serves as a selective probe for post-assembly trafficking and egress steps.

The M protein is an essential structural component of the virion that forms the scaffold for envelope formation and interacts directly with the other structural proteins for their incorporation during assembly. Previous studies showed that subtle changes in the carboxy end of the M proteins are detrimental or negatively impact virus assembly (31, 32). It was also shown that virus-like-particles (VLPs) were not assembled when the monomeric red fluorescent protein (mRFP1) tag was added to the carboxy terminus of the SARS M protein (33). A recent report showed that fluorescently labeled SARS-CoV-2 VLPs are produced only when tagged and untagged M and N proteins are coexpressed (34). We expressed M-GFP in accessory ORF 4 in the MHV genome with WT M expressed in parallel, from its normal ORF. This was necessary since our attempts to replace the WT M gene with tagged M protein in the viral genome were not successful. However, this proved to be useful in that we were able to directly compare the behavior of WT and labeled M in the same cellular milieu. Analysis of immunostained cells fixed at different time points after infection, showed that most M-GFP closely mimics the distribution of native M early in infection, prior to viral assembly. This indicates that M-GFP serves as a reliable proxy for M trafficking during the initial stages of infection. Later in infection, when the majority of WT M is found in the cell periphery, presumably in the form of assembled virions, we observed that much of the M-GFP remained in the cell interior. This observation may help to explain why M-GFP cannot substitute entirely for WT M. M-GFP may not incorporate as efficiently into viral particles and is instead retained at/near the ERGIC.

Recombinant coronaviruses expressing fluorescent tags appended to other CoV structural proteins were previously reported. MHV S-GFP protein was expressed in place of WT S in the viral genome through incorporation by targeted recombination (35). A recombinant MHV with GFP fused to the carboxy end of the N gene was expressed under a transcription regulatory sequence (TRS) between genes 2a and S (36). Even though viruses were recovered, in both cases expression of the chimeric proteins was quickly lost during passage of the viruses. Notably, S and N proteins may traffic within the cell prior to and independently of virus particle assembly (30); thus, S-GFP or N-GFP probes may be less able to distinguish pre- from post-assembly trafficking steps, compared to M-GFP.

Reporter genes have also been expressed in place of CoV accessory ORFs in previous studies. MHV ORF 4, the insertion site of our M-GFP fusion protein, was used previously for expression of single heterologous genes, since it is nonessential (37, 38). Recombinant MHV expressing enhanced GFP (EGFP) was previously selected by targeted recombination for monitoring replication and spread in cell culture and in mice (39). EGFP expression was stable through at least six passages in cell culture, like our results with M-GFP expression. Other CoV ORFs have also been used for heterologous protein expression. GFP expression from ORF3a has been described in a recombinant transmissible gastroenteritis coronavirus (TGEV), and recently from ORF3 in a recombinant swine diarrhea syndrome coronavirus (SADS-CoV) (40) (41, 42). Both were genetically stable for 10 passages. ORF 7 in SARS-CoV-2 was used for expression of GFP, or GFP fused to nanoluciferase, in development of reverse genetic systems for SARS-CoV-2 (37, 43). Expression of heterologous genes from CoV ORFs is clearly useful, but CoVs are apparently under selective pressure to disrupt these heterologous genes, as they are lost over time - especially so for the tagged structural envelope proteins. We observed that M-GFP was lost as the virus was passaged, even though it did not have severe effects on virus replication and spread. It is possible that replacing the ORF 4 coding sequence may not be tolerated well, regardless of what heterologous gene sequences are inserted. We did not explore expression from alternative ORFs.

Co-immunoprecipitations showed that M-GFP and WT M apparently interact, which likely forms the basis for assembly of the fusion protein into particles. The M protein forms a lattice that serves as the scaffold for formation and organization of the viral envelope: cryo-EM tomography showed that M forms local ordered networks in the envelope, and it has been suggested that the protein assumes different long and short conformations during assembly (22, 25). Recently our understanding of M proteins was enhanced with imaging of SARS-CoV-2 M protein in lipid bilayers using cryo-EM and atomic force microscopy (21, 44). Results suggest that M oligomerization facilitates membrane curvature, which is important for assembly of coronavirus pleomorphic round virus particles. How oligomers of M and M-GFP, undergo conformational changes and promote membrane curvature to participate in assembly remains to be determined. We have not yet quantified the ratio of M to M-GFP in virus particles, but our time-course analysis of M and M-GFP co-transport toward the cell surface suggests that more WT M moves to the periphery, whereas more M-GFP is retained in the ERGIC. Since M-GFP is clearly incorporated into particles, we envision that homotypic M-M multimers are likely sufficient to drive the formation of particles, and heterotypic M and M-GFP multimers may come along for the ride. Including a flexible linker peptide between M and GFP in our fusion may allow the M carboxy terminus to assume its normal function, with the GFP extending into the intravirion space. It appears that homotypic M-GFP multimers do not support assembly since we were unable to rescue a virus expressing only M-GFP. Regardless, M-GFP incorporation into virus particles opens new opportunities for tracking and structural studies, as well as replication and pathogenesis.

To address the utility of the M-GFP virus for assessing mechanisms of transport and egress from cells, TIRF imaging in live cells was explored. This modality excludes most of the cell volume from view and allowed us to image only the M-GFP that was successfully transported from the bottom (adherent) cell surface. In each infected cell, we were able to detect and track up to several thousand GFP-positive particles moving near the plasma membrane, over the course of two minutes of imaging. The approximate velocity of M-GFP particles at ∼1.67 um/s is within the range normally reported for intracellular vesicular transport of cellular cargoes (28, 29). Although particles tracked for long duration were, necessarily, those with a trajectory primarily parallel to the membrane, we occasionally observed a particle that would suddenly appear with high intensity and subsequently remain fixed in place. This behavior is very similar to exocytosis events observed during herpesvirus exocytosis (45-48). Further studies are needed to confirm whether the observed phenomenon represents exocytosis of MHV particles.

The size of the fluorescent puncta observed by TIRF was slightly less than 500 nm, which is much larger than the size of roughly 100 nm coronavirus particles. This is consistent with the diffraction limit of light microscopy but based on transmission electron microscopy studies larger secretory organelles may carry many virus particles to the plasma membrane for bulk release (49). However, we noted that there was very little variation in the observed size of the fluorescent puncta; therefore, most viral secretory organelles are likely below the diffraction limit.

Our results show that the M-GFP recombinant virus is a useful tool for probing details of coronavirus assembly and egress in live cells during infection. Previous studies have identified the intracellular compartments that viral proteins traffic through during assembly, and recent cryo-EM tomography studies provide further insights (5, 49, 50). However, there is still much to be learned about the details of coronavirus assembly and trafficking as virions exit from infected cells. For example, there is limited information about the host proteins and cellular pathways that directly interact with viral components and cargo transport vesicles during virus egress. Recently it was reported that β-CoVs use the lysosome-related organelles for egress, rather than post-Golgi secretory carriers as described previously (51-53). The microtubule- and actin-based motors involved in CoVs intracellular transport, regardless of the egress pathway, are not known. We recently reported, based on a yeast two-hybrid screen and transient expression in mammalian cells, that MHV-CoV M protein tail interacts with a splice variant of myosin Vb, and that trafficking may involve Rab10 (54). This prompted us to make the recombinant M-GFP virus to gain insight and explore the significance of these interactions and others in the context of virus infection. Recovery of the M-GFP virus has expanded our toolset for addressing the potential role of myosin or other motors, host protein trafficking regulators and pathways involved in CoV assembly and egress. Labeled virus particles may also be used for study of other processes during CoV infection, including possibly viral entry, replication steps in close parallel with particle assembly and the contributions of these processes to pathological changes during infection.

## MATERIALS AND METHODS

### Cells and viruses

HEK 293T (ATCC), mouse L2 and 17 clone1 (17Cl1) cells were maintained in Dulbecco’s modified Eagle’s medium (DMEM) containing 10% heat-inactivated fetal calf serum (FCS), plus penicillin, streptomycin, and L-glutamine. Virus stocks were grown in mouse 17Cl1 or L2 cells. Virus titers were determined in L2 cells. Baby hamster kidney (BHK) cells expressing the MHV carcinoembryonic antigen-related cell adhesion molecule 1 (CEACAM1) receptor were used for recovery of M-GFP virus (55).

### Reverse genetics

The M-GFP virus was made by reverse genetics using a MHV A59 infectious clone (55). The M-GFP fusion coding sequence was engineered in place of the ORF4 coding region in the G-clone that encompasses ∼8.7 kb of the 3’ end of the genome which includes the structural genes (55). A cDNA PCR product containing the M-GFP fusion protein coding sequence replaced the coding region between the SbfI and EcoRV sites in the G-clone. Full-length cDNA clones were assembled, transcribed, and electroporated into BHK-MHVR cells basically as we described previously (56). Briefly, following electroporation BHK-MHVR cells were seeded concurrently with L2 cells. At 72 h after electroporation media were harvested and an aliquot was used to infect L2 cells. The medium was removed from infected cells at ∼24 hpi. Total cytoplasmic RNA from cells remaining on the flasks was extracted using TRIzol Reagent (Invitrogen). RNA was reverse transcribed using the Superscript IV RT-PCR system (Invitrogen) using gene specific primers an oligo (dT) primer. After first strand synthesis the RT product was subjected to 20 cycles of PCR amplification using KAPA HiFi hot start polymerase (Roche) and appropriate primers to amplify the sequence from the end of the S gene to the end of the untranslated region (UTR). PCR products were cleaned with QIAquick PCR purification (Qiagen) before being sequenced directly.

Viruses were subsequently serially plaque purified from media taken directly off cells that had been electroporated. Multiple plaque isolates were passaged on 17Cl1 to generate virus stocks. RNA was extracted from L2 infected cells following passage of plaque purified viruses for RT-PCR and sequence confirmation of the the 3’ end of the genome.

### Virus Growth Kinetics

Growth kinetic experiments were carried out in monolayers of mouse 17Cl1 cells, which were infected with either WT MHV or M-GFP plaque purified virus at a multiplicity of infection (MOI) of 0.01 PFU/cell essentially as previously described (56). Cell culture supernatants were collected at the indicated time points after infection and titers were determine by plaque assay on mouse L2 cells. Low-melt agarose-media overlays (1:1 mix of 2X MEM containing 4% FBS and 3% low agarose) were removed at 72-96 hpi. Cells were fixed and stained with 0.2% (w/v) crystal violet in 20% ethanol to visualize plaques. Three independent plaque assay experiments were performed in duplicate or triplicate in 6-well plates. The number of plaques were averaged for the three experiments. The significance between the WT and M-GFP virus titers from the 48 hpi was determine using Welch’s t-test (Student’s t-test for unequal variances).

### Immunoprecipitation (IP)

Monolayers of L2 cells were infected with WT MHV or M-GFP virus stock at a MOI of 1. After adsorption for 1 h, cells were washed with phosphate buffered saline (PBS) before feeding with Dulbecco’s Modified Eagle’s Medium (DMEM) containing 2% heat inactivated fetal bovine serum (FBS). At 24 hpi cells and media were separated by centrifugation. The extracellular media was stored at -80°C until further processing. Pelleted cells were lysed on ice for 30 min with 500 µl of RIPA buffer (1% Triton X-100, 1% deoxycholate, 150 mM NaCl, 50 mM Tris-HCl [pH7.6], 20 mM EDTA) plus 0.1% SDS and cOmplete ULTRA (Sigma) protease inhibitor. One hundred µl of the post-nuclear lysate was reserved for western blotting. The SDS concentration for the remaining 400 μl was increased to 0.3%. Half of the lysate was incubated overnight at 4° with anti-M monoclonal antibody J1.3 that recognizes the amino end of the protein (57). The other half of the sample was incubated in parallel with anti-GFP monoclonal antibody GF28R (Invitrogen). Antigen-antibody complexes were isolated by incubation with protein A-Sepharose for 2h at 4°C with rocking. Pelleted complexes were washed with RIPA buffer containing detergent followed by a final wash without detergent before elution by incubation for 5 min at 37°C in Laemmli SDS-PAGE sample loading containing 10% β-mercaptoethanol (βME). Half of each sample was analyzed by SDS-PAGE and western blotting.

Extracellular fractions were ultracentrifuged over a 20% sucrose cushion for 3 h using a Sorvall SW 55Ti rotor at 4°C). Pelleted virus was re-suspended in 1 ml of RIPA buffer (100 µl of TMEN buffer pH 6.0 (50 mM Tris-HCl, 50 mM maleic acid, 1 mM EDTA, 100 mM NaCl). One hundred µl was set aside for the input control and the remaining two 450 µl aliquots were used for IP, following the same protocol described above for the intracellular fraction.

### Western Blotting

Cells were lysed on ice in buffer containing 150 mM NaCl, 0.1% Triton X-100, 50 mM Tris-HCl (pH 8), and 1mM of phenylmethylsulphonyl fluoride (PMSF) for 30 min. Lysates were centrifuged at 13,500 x g for 3 min. Post-nuclear lysates were mixed with 2X Laemmli SDS-PAGE sample buffer plus βME for SDS-PAGE. Proteins were transferred to a PVDF membrane and analyzed with the anti-MHV M 9246 that recognizes the carboxy terminal 18 amino acids of the protein that was made in the B. Hogue lab and polyclonal anti-GFP A6455 (Thermo Fisher Scientific) described above, followed by appropriate secondary antibodies.

### Fluorescence Microscopy

For imaging of purified virus, collected fractions were diluted 1:5 with PBS and approximately 20 µL of suspension was dropped onto a #1.5 cover glass and imaged immediately. Images were collected for M-GFP and wild-type control virus on a Nikon AX R confocal, using a 60x, 1.4 N.A. lens and 488 nm laser excitation. For immunofluorescence, cells grown in an 8-well chambered coverglass (Ibidi) were fixed at the indicated time after infections in 4% formaldehyde in PBS for 5 minutes, permeabilized for 90 seconds with Karsenti’s lysis buffer (0.5% Triton X-100, 80 mM PIPES, 1.0 mM MgSO_4_, 5.0 mM EGTA, pH 7.0), then fixed for an additional 2 min in 4% formaldehyde. Cells were rinsed three times in PBS then blocked with 5% goat serum in PBS for 30 minutes. Primary antibodies were diluted in 1% goat serum in PBST (PBS plus 0.05% Tween 20, pH 7.4). Mouse, anti-MHV M J1.3 monoclonal (26) was used 1:500, rabbit, anti-ERGIC-53 (Proteintech, Catalog # 13364-1-AP) was used to probe the ERGIC compartment at 1:300 dilution. Cells were incubated overnight at 4°C in primary antibody then rinsed three times in PBST. Secondary antibodies, Alexa Fluor 568 anti-mouse (ThermoFisher Scientific, Catalog # A-11004) and Alexa Fluor 633 anti-rabbit (ThermoFisher Scientific, Catalog # A-21070) were diluted 1:1000 and samples were incubated for 1 hour at room temperature with gentle rocking. Samples were rinsed three time in PBST, then once in PBS. Finally, nuclei were labeled with DAPI (1 µg/mL in PBS). Images were collected with a Nikon AX R confocal, using a 60x, oil immersion, 1.4 N.A. lens. For live cell TIRF imaging, cells were grown in a chambered cover glass. At the indicated time after infection, samples were imaged by TIRF on a Nikon Ti-2 microscope base equipped with a Tokai-Hit stage-top incubator maintained at 37°C with 5% CO_2_. A 488 nm laser and a 60x, 1.49 N.A TIRF lens were used for TIRF excitation. Cells were imaged for 1.5-2 minutes per field. For fixed TIRF imaging, 17C1 cells were fixed at 6.5 hpi and immunolabed as described above for WT M. S protein was detected using goat anti-MHV S primary antibody (30) diluted 1:250 and Alexa Fluor and AlexaFluor Plus 405, anti-goat secondary antibody (ThermoFisher Scientific, Catalog # A48259) as described above. Three color TIRF images were acquired simultaneously using 405, 488, and 561 nm lasers to image S, M-GFP, and WT M, respectively.

Mouse L2 cells were infected with M-GFP virus at a MOI of 1 for passage stability analysis. At 24 hpi cells were fixed, permeabilized and blocked as described above. Cells were incubated overnight at 4℃ with anti-M polyclonal primary antibody 9246. Cells were washed and further incubated at room temperature with Alexa Fluor 568 anti-rabbit secondary antibody (ThermoFisher Scientific). Slides were rinsed before staining nuclei with DAPI. Images were collected using a Nikon AX R confocal microscopy using a 60X oil immersion, 1.4 N.A. lens.

### Electron Microscopy

Gradient-purified virus particles were deposited on Formvar/carbon film 10/3-4 nm on square 300 mesh copper (FCF300-Cu-SB) grids and negatively stained with 2.0% uranyl acetate in water. Virus particles were imaged using a Talos L120C TEM (ThermoFisher Scientific). For cryo-electron microscopy (cryo-EM), gradient-purified virus particles were deposited on copper C-flat holey carbon grids and vitrified by plunge-freezing with liquid ethane (Vitrobot). Grids were stored in liquid nitrogen until imaged using a Titan Krios 80/300 KeV cryo-TEM/STEM (FEI).

### Image Analysis

For colocalization experiments, an analysis pipeline was constructed in Nikon Elements General Analysis 3 (GA3) software. First, images were denoised using the Denoise AI module, then background subtraction was done by the rolling ball method with a radius of 4 µm. A threshold was applied to each channel automatically using the Otsu method to generate binary overlays. Regions of interest were defined by combining pairs of binary masks by an “Or” function to determine areas positive for either of the signals to be compared. For example, the binary mask for the green signal from the M-GFP was combined with the binary mask by a Boolean OR function to determine areas positive for ether of the signals to be compared. For example, the binary mask for the red signal from the WT M fluorescence to define the area to be assessed for colocalization of those species. Mander’s Correlation Coefficients (MCC) were calculated to determine the fractional overlap of signal 1 with signal 2 (M1) and signal 2 with signal 1(M2). This was done to determine the fraction of M-GFP signal that was coincident with WT M signal (M1) and the fraction of WT M signal that was coincident with M-GFP signal (M2). M1 was also calculated in the same fashion to determine the fraction of either WT M or M-GFP within the ER-Golgi Intermediate Compartment (ERGIC). We determined the intensity correlation between the WT M and M-GFP signals by calculating the Pearson’s Correlation Coefficient (PCC). This was done by defining the areas of interest by combining binaries in the same way as described above. However, the input for the PCC was the original, background subtracted, channels with the intensity values. Statistical significance was determined by ANOVA using Graphpad Prism 10.2.3. For particle tracking in live cell TIRF movies, data were preprocessed by denoising, rollingball background subtraction, and Otsu thresholding to generate binary images as described above. The Nikon Elements tracking module was then used to measure the motility of particles. Particles were included in the analysis if they had a line length (total x-y displacement) of greater than 0.5 µm, and a line speed greater than 0.01 µm/s. We also limited analysis to particles with speed less than 10 µm/s. This eliminated particles that had been released from cells which could be seen diffusing rapidly in the medium.

## ACKNOWLEDGMENTS

This work was supported by NIH NIDDK subaward VUMC95054 (BGH) under parent grant R01DK048370 (JRG), NSF STC Award 1231306 (BGH), NSF RAPID: IIBR Award 2032199 (BGH). Partial support was provided by More Graduate Education at Mountain States Alliance (MGE@MSA) Alliance for Graduate Education and the Professoriate (AGEP) National Science Foundation (NSF) Cooperative Agreement No. HRD-997886 (BS) and NIH NINDS R01NS117513 (IBH).

We thank Dr. Dewight Williams for help with cryoEM imaging, and School of Life Sciences Undergraduate Research Program (SOLUR) student Lindsey (Ngoc Xuan Nhi) Cai for help with construction of the recombinant M-GFP virus. Microscopy was conducted in the Biodesign Imaging Facility, Advanced Light Microscopy Core and the Eyring Materials Center, John M. Cowley Center for High-Resolution Electron Microscopy, ASU Knowledge Enterprise Core Facilities.

## Author contributions

B.S.: conceptualization, experiments, data collection and analysis, methodology, review and editing. N.G.: experiments, data collection and analysis, methodology, writing, editing, review. L.A.L. and J.R.G.: data analysis and review.

I.B.H.: data analysis, writing, editing, review. H.G. conceptualization, experimental design, experiments, data collection and analysis, writing, editing. B.G.H: conceptualization, experimental design and data analysis, writing, and editing.

## FIGURE LEGENDS SUPPLEMENT

**Fig. S1.**
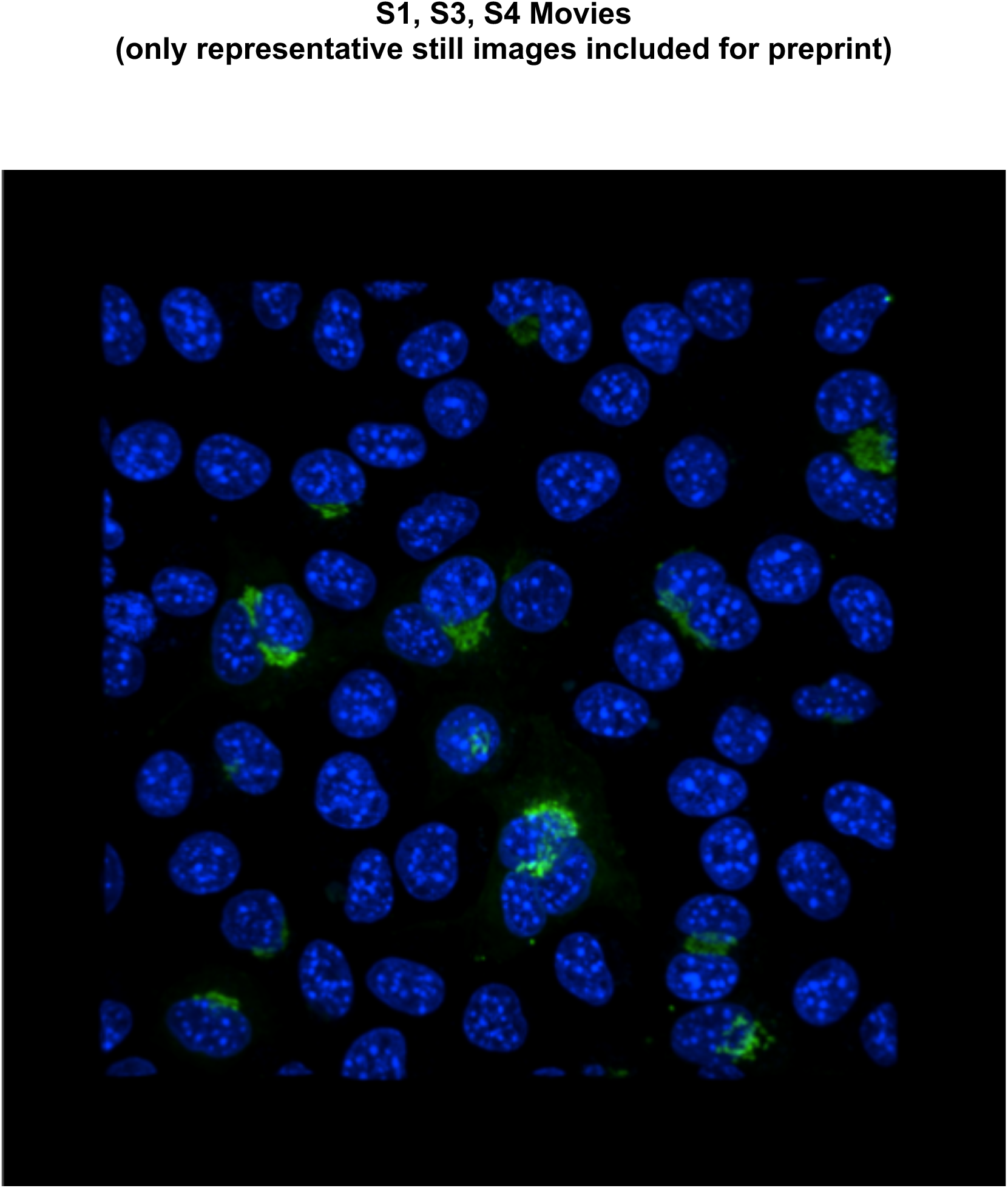
M-GFP used to track the progression of MHV infection in live cells. L2 cells were infected with M-GFP MHV. Nuclei were labeled with Hoechst. Cells were imaged by confocal. Z-stacks were collected every 2 minutes from 5-10 hpi. Movie is a 3D rendered view from the perspective of looking down on the cells from above. Green=GFP; blue=nuclei (Hoechst).

**Fig. S2.**
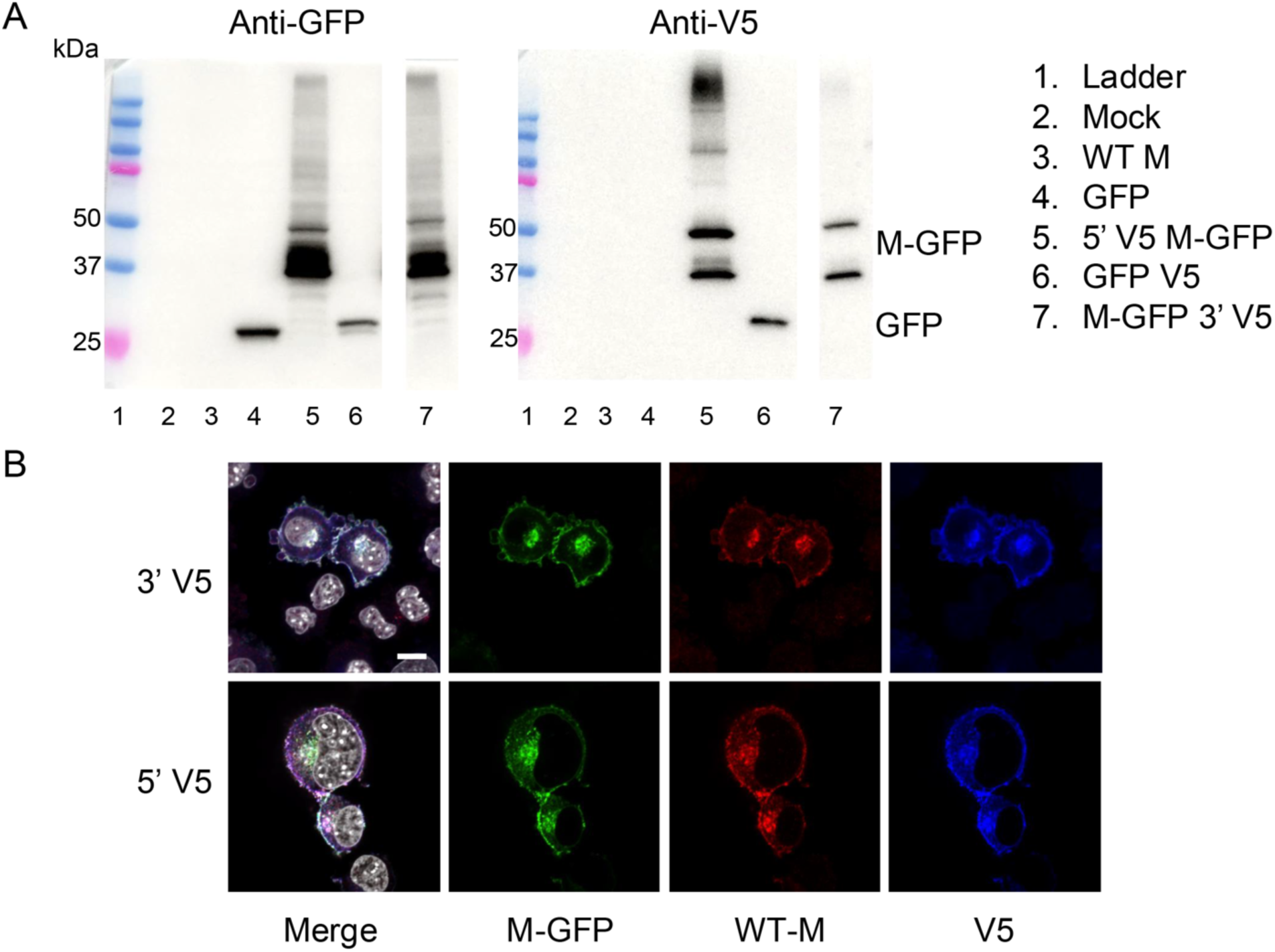
M-GFP protein remains intact when expressed. (A) L2 cells were transfected with untagged WT M or GFP; M-GFP with a V5 tag at the amino end; GFP or M-GFP with a V5 tag at the carboxy end. At 24 h after transfection cell lysates were immunoblotted and probed with antibodies against GFP or V5. When probed with anti-GFP and anti-V5, the 5’ V5 M-GFP and the 3’ V5 M-GFP show bands 37kDa and 50 kDa (lanes 5 and 7). (B) L2 cells were transfected with M-GFP with the amino V5 tag (top row) or amino V5 tag (bottom row) (B) At 24 hours after transfection, cells were immunolabeled for WT M and V5, imaged by confocal. Typical fields are shown selected from at least 3 fields acquired from each of 2 independent experiments. White=nuclei (DAPI); green=M-GFP; red=WT M; blue=V5. Scale bar = 10 µm.

**Fig. S3.**
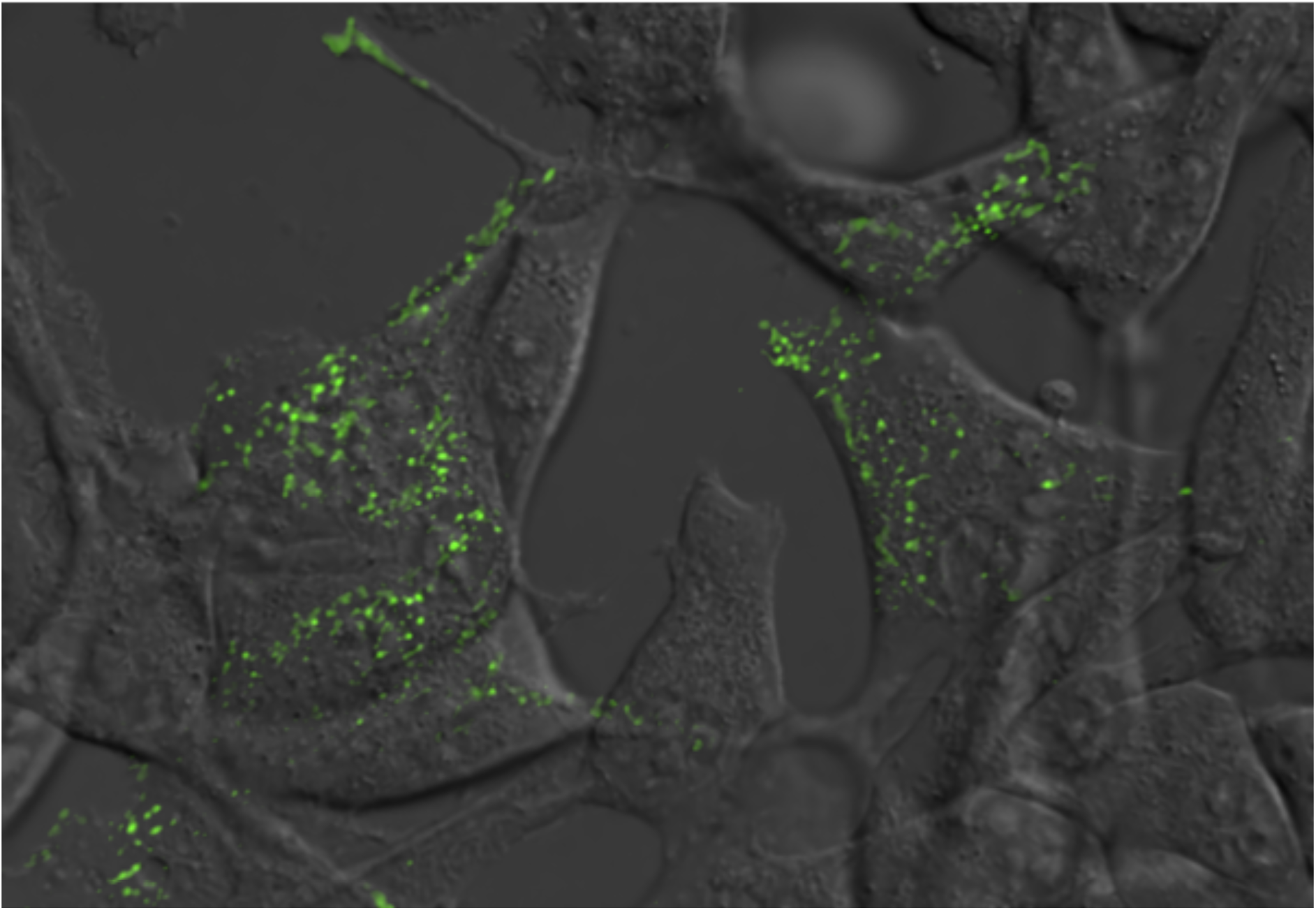
M-GFP allows imaging of assembled MHV virus trafficking at and near the plasma membrane. 17C1 cells were infected with M-GFP MHV then imaged live by TIRF microscopy at 6-7hpi. TIRF was collected in the GFP channel and overlaid on differential interference contrast, still image. Timelapse images were collected for up to 2.5 minutes per field. A typical timelapse is shown, selected from at least 5 datasets taken from each of 3 independent experiments.

**Fig. S4.**
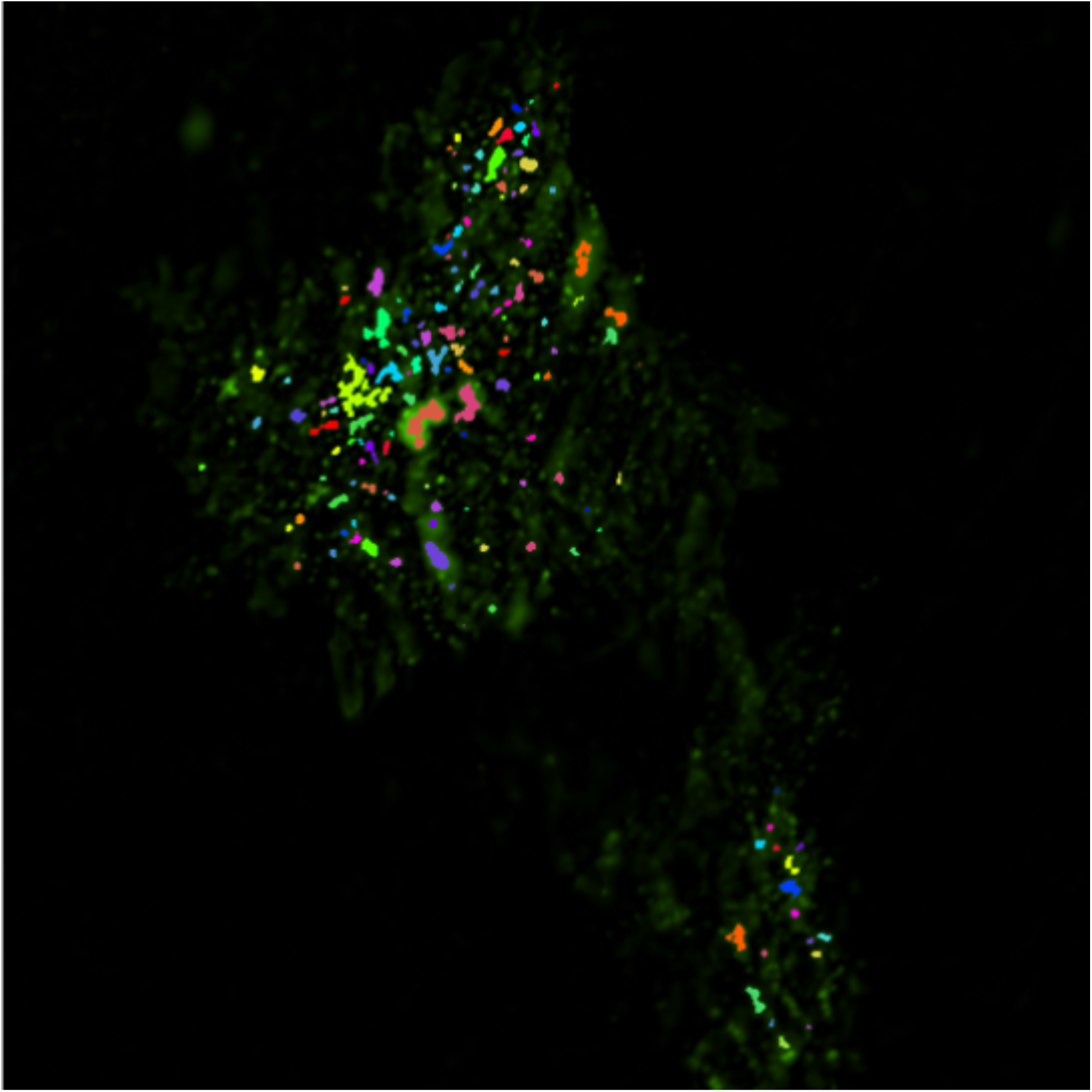
Tracking of virus at and near the plasma membrane in 17Cl1 infected cells. Timelapse images collected by TIRF (as described in Fig. S3) were analyzed by Nikon Elements particle tracking module. Movie shows tracks detected from 60 seconds of data collection.

## SUPPLEMENTAL MATERIALS AND METHODS

### Microscopy

For live cell z-stack, time-lapse images, L2 cells were grown in an 8-well Ibidi chambered coverglass. After infection with M-GFP MHV, nuclei were labeled with Hoechst. Images were collected on a Nikon AX R confocal, using a 60x, oil immersion, 1.4 N.A. lens. The Nikon Ti-2 microscope base was equipped with a Tokai-Hit stage-top incubator which was maintained at 37°C and 5% CO2. Cells were imaged from 5-10 hpi. Full z-stacks through the cells were collected every 2 minutes. Fixed immunofluorescent imaging, live cell TIRF, and particle tracking were done as described in main text.

## Notes

Authors declare no conflicts of interest.

### Competing Interest Statement

The authors have declared no competing interest.

